# Revisiting the links-species scaling relationship in food webs

**DOI:** 10.1101/2020.02.13.947531

**Authors:** Andrew MacDonald, Francis Banville, Timothée Poisot

## Abstract

Predicting the number of interactions that species in a food web will establish is an important task. These trophic interactions underlie many ecological and evolutionary processes, ranging from biomass fluxes, ecosystem stability, resilience to extinction, and resistance against novel species. We investigate and compare several ways to predict the number of interactions in food webs. We conclude that a simple beta-binomial model outperforms other models, with the added desirable property of respecting biological constraints. We show how this simple relationship gives rise to a predicted distribution of several quantities related to link number in food webs, including the scaling of network structure with space, and the probability that a network will be stable.

## Introduction

Community ecologists are fascinated by counting things. It is therefore no surprise that early food web research paid so much attention to counting species, counting trophic links, and uncovering the relationship that binds them – and it is undeniable that these inquiries kickstarted what is now one of the most rapidly growing fields of ecology [1]. More species (*S*) always means more links (*L*); this scaling is universal and appears both in observed food webs and under purely neutral models of food web structure [2]. In fact, these numbers underlie most measures used to describe food webs [3]. The structure of a food web, in turn, is almost always required to understand how the community functions, develops, and responds to changes [4,5], to the point where some authors suggested that describing food webs was a necessity for community ecology [6,7]. To this end, a first step is to come up with an estimate for the number of existing trophic links, through sampling or otherwise. Although both *L* and *S* can be counted in nature, the measurement of links is orders of magnitude more difficult than the observation of species [8,9]. As a result, we have far more information about values of *S*. In fact, the distribution of species richness across the world is probably the most frequently observed and modelled ecological phenomena. Therefore, if we can predict *L* from *S* in an ecologically realistic way, we will be in a position to make first order approximations of food web structure at large scales, even under our current data-limited regime.

Measures of food web structure react most strongly to a handful of important quantities. The first and most straightforward is *L*, the number of trophic links among species. This quantity can be large,especiallyinspecies-richhabitats,butitcannotbearbitrarilylarge. It is clear to any observer of nature that of all imaginable trophic links, only a fraction actually occur. If an ecological community contains *S* species, then the maximum number of links in its food web is *S*^2^: a community of omnivorous cannibals. This leads to the second quantity: a ratio called *connectance* and defined by ecologists as *Co* = *L/S*^2^. Connectance has become a fundamental quantity for nearly all other measures of food web structure and dynamics [10]. The third important quantity is another ratio: *linkage density*, *L_D_* = *L/S*. This value represents the number of links added to the network for every additional species in the ecological system. A closely related quantity is *L_D_* × 2, which is the *average degree*: the average number of species with which any taxa is expected to interact, either as predator or prey. These quantities capture ecologically important aspects of a network, and all can be derived from the observation or prediction of *L* links among *S* species.

Because *L* represents such a fundamental quantity, many predictive models have been considered over the years. Here we describe three popular approaches before describing our own proposed model. The *link-species scaling(LSSL)* [11] assumes that all networks have the same *average degree*; that is, most species should have the same number of links. Links are modelled as the number of species times a constant:

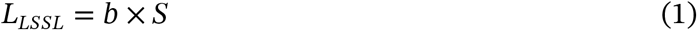

with *b* ≈ 2. This model imagines that every species added to a community increases the number of links by two – for example, an animal which consumes one resource and is consumed by one predator. This model started to show its deficiencies when data on larger food webs became available: in these larger webs, *L* increased faster than a linear function of *S*. Perhaps then all networks have the same *connectance* [12]? In other words, a food web is always equally filled, regardless of whether it has 5 or 5000 species. Under the so-called “constant connectance” model, the number of links is proportional to the richness squared,

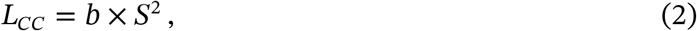

where *b* is a constant in] 0,1 [representing the expected value of connectance. The assumption of a scaling exponent of 2 can be relaxed [12], so that *L* is not in direct proportion to the maximum number of links:

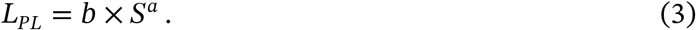

This “powerlaw” model can be parameterized in many ways, including spatial scaling and species area relationships [13]. It is also a general case of the previous two models, encompassing both link-species scaling (*a* = 1, *b* ≈ 2) and the strict constant connectance (*a* = 2,0 < *b* < 1) depending on which parameters are fixed. Power laws are very flexible, and indeed this function matches empirical data well – so well that it is often treated as a “true” model which captures the scaling of link number with species richness [14–16], and from which we should draw ecological inferences about what shapes food webs. However, this approach is limited, because the parameters of a power law relationship can arise from many mechanisms, and are difficult to reason about ecologically.

But the question of how informative parameters of a power law can be is moot. Indeed, both the general model and its variants share an important shortcoming: they cannot be used for prediction while remaining within the bounds set by ecological principles. This has two causes. First, models that are variations of *L* ≈ *b* × *S^a^* have no constraints - with the exception of the “constant connectance” model, in which *L*_cc_ has a maximum value of *S*^2^. However, we know that the number of links within a food web is both lower and upper bounded [12,17]: there can be no more than S^2^ links, and there can be no fewer than *S* — 1 links. This minimum of *S* — 1 represents communities where all species interact and at least some of the organisms are heterotrophs [12]. Numerous simple food webs could have this minimal number of links – for example, a linear food chain wherein each trophic level consists of a single species, each of which consumes only the species below it; or a grazing herbivore which feeds on every plant in a field. Thus the number of links is constrained by ecological principles to be between *S*^2^ and *S* − 1, something which no present model includes. Secondly, accurate predictions of *L* from *S* are often difficult because of how parameters are estimated. This is usually done using a Gaussian likelihood for *L*, often after log transformation of both *L* and *S*. While this approach ensures that predicted values of *L* are always positive, it does nothing to ensure that they stay below *S*^2^ and above *S* − 1. Thus a good model for *L* should meet these two needs: a bounded expression for the average number of links, as well as a bounded distribution for its likelihood.

Here we suggest a new perspective for a model of *L* as a function of *S* which respects ecological bounds, and has a bounded distribution of the likelihood. We include the minimum constraint by modelling not the total number of links, but the number in excess of the minimum. We include the maximum constraint in a similar fashion to the constant connectance model described above, by modelling the proportion of flexible links which are realized in a community.

## Interlude - deriving a process-based model for the number of links

Based on the ecological constraints discussed earlier, we know that the number of links *L* is an integer such that *S* − 1 ≤ *L* ≤ *S*^2^. Because we know that there are at least *S* − 1 links, there can be at most *S*^2^ − (*S* − 1) links *in excess* of this quantity. The *S* − 1 minimum links do not need to be modelled, because their existence is guaranteed as a pre-condition of observing the network. The question our model should address is therefore, how many of these *S*^2^ − (*S* − 1) “flexible” links are actually present? A second key piece of information is that the presence of a link can be viewed as the outcome of a discrete stochastic event, of which the alternative outcome is that the link is absent. We assume that all of these flexible links have the same chance of being realized, which we call *p*. Then, if we aggregate across all possible species pairs, the expected number of links is

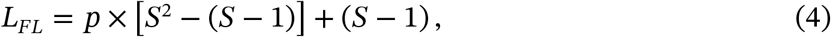

where *p* ∈ [0,1]. When *p* = 1, *L* is at its maximum (*S*^2^), and when *p* = 0 it is at the minimum value (*S* − 1). We use the notation *L_FL_* to represent that our model considers the number of “flexible” links in a food web; that is, the number of links in excess of the minimumbutbelow the maximum. Because we assume that every flexible link is an independent stochastic event with only two outcomes, we can follow recent literature on probabilistic ecological networks [18] and represent them as independent Bernoulli trials with a probability of success p. Furthermore, the observation of *L* links in a food web represents an aggregation of S^2^ − (*S* − 1) such trials. If we then assume that *p* is a constant for all links in a particular food web, but may vary between food webs (a strong assumption which we later show is actually more stringent than what data suggest), we can model the distribution of links directly as a shifted beta-binomial variable:

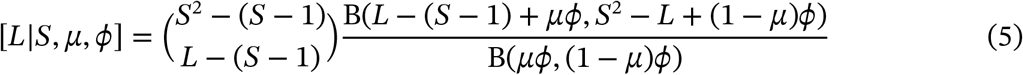

Where B is the beta function, *μ* is the average probability of a flexible link being realized (*i.e*. the average value of *p* across networks in the dataset) and *ϕ* is the concentration around this value. The support of this distribution is limited to only ecologically realistic values of *L*: it has no probability mass below *S* − 1 or above *S*^2^. This means that the problem of estimating values for μ and φ is reduced to fitting the univariate distribution described in eq. (12). For more detailed explanation of the model derivation, fitting, and comparison, see Experimental Procedures.

In this paper we will compare our flexible links model to three previous models for *L*. We estimate parameters and compare the performance of all models using open data from the `mangal.io` networks database [19]. We show how this model not only outperforms existing efforts at predicting the number of links, but also has numerous desirable properties from which novel insights about the structure of food webs can be derived.

## Results and Discussion

### Flexible links model fits better and makes a plausible range of predictions

All models fit well, without any problematic warnings (see Experimental Procedures), and our model for flexible links outperformed previous solutions to the problem of modelling *L*. The flexible links model, which we fit via a beta-binomial observation model, had the most favourable values of PSIS-LOO information criterion (table 1) and of expected log predictive density (ELPD), relative to the three competing models which used a negative binomial observation model. Pareto-smoothed important sampling serves as a guide to model selection [20]; like other information criteria it approximates the error in cross-validation predictions. Smaller values indicate a model which makes better predictions. The calculation of PSIS-LOO can also provide some clues about potential model fits; in our case the algorithm suggested that the constant connectance model was sensitive to extreme observations. The expected log predictive density (ELPD), on the other hand, measures the predictive performance of the model; here, higher values indicate more reliable predictions [20]. This suggests that the flexible links model will make the best predictions of *L*.

**Table 1:** Comparison of the four different models. We show Pareto-smoothed important sampling values (PSIS-LOO) and their standard deviation. PSIS-LOO is similar to information critera in that smaller values indicate better predictive performance. We also show expected log predictive density (ELPD) differences to the maximum for all models, along with the standard error (SE) of these differences.

| Model | eq. | PSIS-LOO | ΔELPD | $SE_{\Delta$ELPD |
| --- | --- | --- | --- | --- |
| Flexible links | 4 | $2520.5 \pm 44.4$ | 0 | 0 |
| Power law [13] | 3 | $2564.3 \pm 46.6$ | -21.9 | 6.5 |
| Constant [12] | 2 | $2811.0 \pm 68.3$ | -145.3 | 21.1 |
| Link-species scaling [11] | 1 | $39840.1 \pm 2795.1$ | -18659.8 | 1381.7 |

To be useful to ecologists, predictions of *L* must stay within realistic boundaries determined by ecological principles. We generated posterior predictions for all models and visualized them against these constraints (fig. 1). The LSSL model underestimates the number of links, especially in large networks: its predictions were frequently lower than the minimum *S* − 1. The constant connectance and power lawmodels also made predictions below this value, especially for small values of *S*. The flexible link model made roughly the same predictions, but within ecologically realistic values.

**Figure 1:**
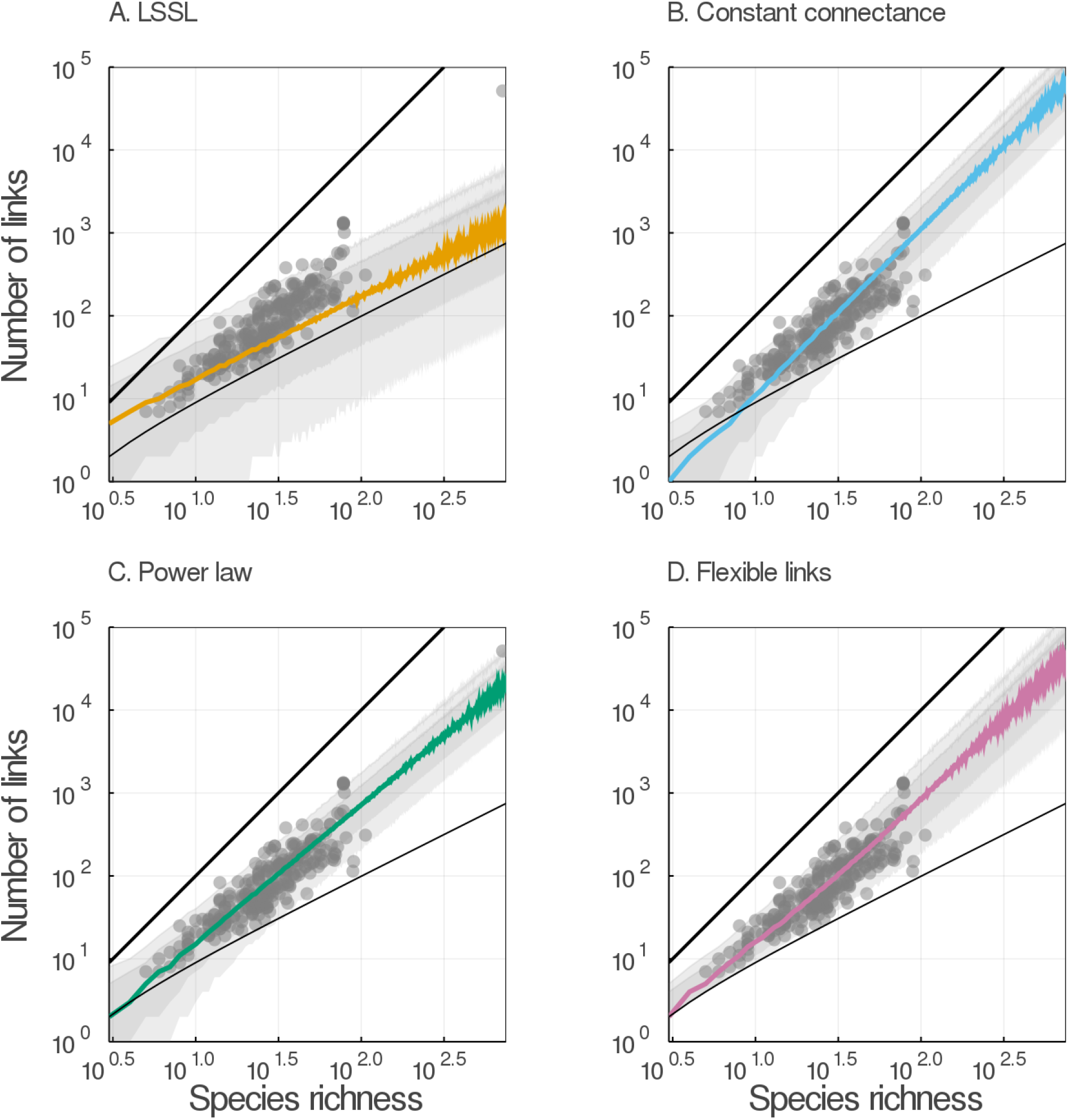
The flexible links model fits better and makes a plausible range of predictions. The number of links is plotted as a function of species richness obtained from the posterior distributions of A) the link-species scaling, B) the constant connectance, C) the power law and D) the flexible links models. In each panel, the colored line represent the median predicted link number and the grey areas cover the 78%and 97%percentile intervals. Empirical datafrom the `mangal.io` database are plotted in each panel (grey dots), as well as the minimal *S* − 1 and maximal *S*^2^ number of links (thinner and bolder black lines, respectively).

### The flexible links model makes realistic predictions for small communities

Constraints on food web structure are especially important for small communities. This is emphasized in fig. 2, which shows that all models other than the flexible links model fail to stay within realistic ecological constraints when *S* is small. The link-species scaling model made around 29% of unrealistic predictions of link numbers for every value of *S* (3 ≤ *S* ≤ 750). The constant connectance and power law models, on the other hand, also produced unrealistic results but for small networks only: more than 20% were unrealistic for networks comprising less than 12 and 7 species, respectively. Only the flexible links model, by design, never failed to predict numbers of links between *S* − 1 and *S*^2^. It must be noted that unrealistic predictions are most common in the shaded area of fig. 2, which represents 90% of the empirical data we used to fit the model; therefore it matters little that models agree for large *S*, since there are virtually no such networks observed.

**Figure 2:**
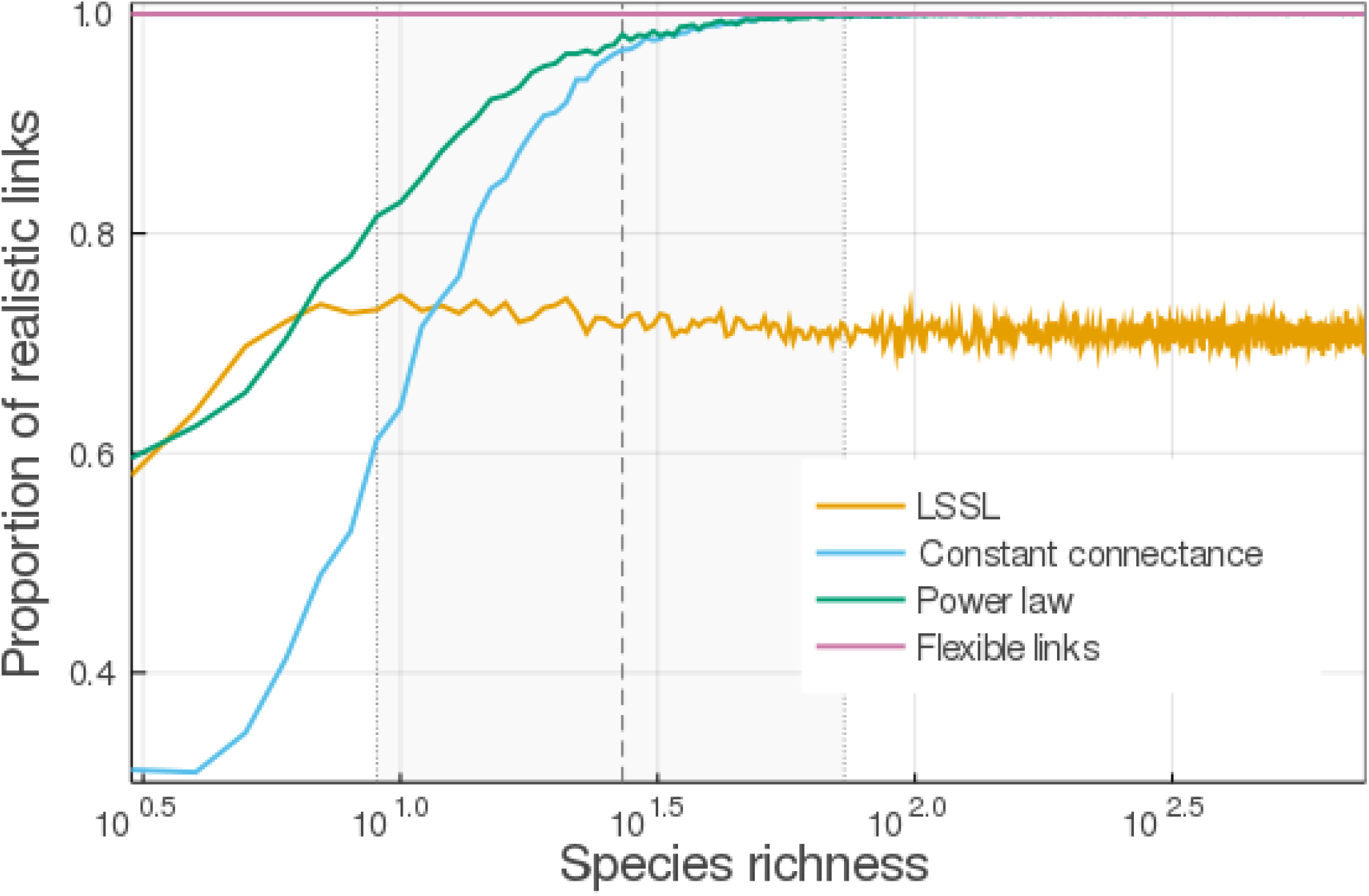
Only the flexible links model makes realistic predictions for small communities. Here we show the proportion of posterior predictions from each of our 4 models which fall outside ecologically realistic values. The proportion of predictions in the correct range increases with species richness for the constant connectance and power law models. Shaded area shows the 5%, 50% and 95% quantiles of the distribution of *S*, demonstrating that many communities have potentially incorrect predictions under previous models.

### Parameter estimates for all models

Although we did not use the same approach to parameter estimation as previous authors, our approach to fitting these models recovered parameter estimates that are broadly congruent with previous work. We found a value of 2.2 for *b* of the LSSL model (table 2), which is close to the original value of approximately 2 [11]. Similarly, we found a value of 0.12 for *b* of the constant connectance model, which was consistent with original estimate of 0.14 [12]. Finally, the parameter values we found for the power law were also comparable to earlier estimates [13]. All of these models were fit with a negative binomial observation model, which has an additional parameter, *κ*, which is sometimes called a “concentration” parameter. This value increases from the top of our table to the bottom, in the same sequence as predictive performance improves in table 1. This indicates that the model predictions are more concentrated around the mean predicted by the model (table 2, column 1).

**Table 2:** Parameter estimates for all models. Mean and standard deviation (SD) is given for each parameter.

| Model | parameter | interpretation | value | SD |
| --- | --- | --- | --- | --- |
| $bS$ [11] | $b$ | links per species | 2.2 | 0.047 |
| | $\kappa$ | concentration of $L$ around mean | 1.4 | 0.12 |
| $bS^2$ [12] | $b$ | proportion of links realized | 0.12 | 0.0041 |
| | $\kappa$ | concentration of $L$ around mean | 4.0 | 0.37 |
| $bS^a$ [13] | $b$ | proportion of relationship | 0.37 | 0.054 |
| | $a$ | scaling of relationship | 1.7 | 0.043 |
| | $\kappa$ | concentration of $L$ around mean | 4.8 | 0.41 |
| $(S^2 - (S - 1))p + S - 1$ | $\mu$ | average value of $p$ | 0.086 | 0.0037 |
| | $\phi$ | concentration around value of $\mu$ | 24.3 | 2.4 |

Our parameter estimates for the flexible links model are ecologically meaningful. For large communities, our model should behave similarly to the constant connectance model and so it is no surprise that *μ* was about 0.09, which is close to our value of 0.12 for constant connectance. In addition, we obtained a rather large value of 24.3 for *ϕ*, which shrinks the variance around the mean of *p* to approximately 0.003 (*var*(*p*) = *μ*(1 − *μ*)/(1 + *ϕ*)). This indicates that food webs are largely similar in their probability of flexible links being realized (thus showing how our previous assumption that *p* might vary between food webs to be more conservative than strictly required). The flexible links model also uses fewer parameters than the power law model and makes slightly better predictions, which accounts for its superior performance in model comparison (table 1)

### Connectance and linkage density can be derived from a model for links

Of the three important quantities which describe networks (*L, Co* and *L_D_*) we have directly modelled *L* only. However, we can use the parameter estimates from our model for *L* to parameterize a distribution for connectance (*L/S*^2^) and linkage density (*L/S*). We can derive this by noticing that eq. (4) can be rearranged to show how *Co* and *L_D_* are linear transformations of *p*:

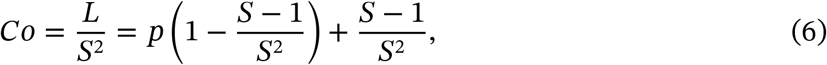

and

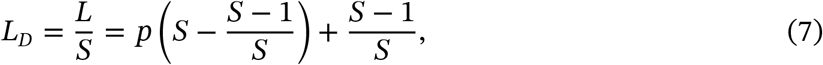

For food webs with many species, these equations simplify: eq. (4) can be expressed as a second degree polynomial, *L_FL_* = *p*×*S*^2^+(1 − *p*)×*S*+(*p* − 1), whose leading term is *p*×*S*^2^. Therefore, when *S* is large, eq. (6) and eq. (7) respectively approach *Co* = *L/S*^2^ ≈ *p* and *L_D_* = *L/S* ≈ *pS*. A study of eq. (6) and eq. (7) also provides insight into the ecological interpretation of the parameters in our equation. For example, eq. (7) implies that adding *n* species should increase the linkage density by approximately *p* × *n*. The addition of 11 new species (*p*^-1^ according to table 2) should increase the linkage density in the food web by roughly 1, meaning that each species in the original network would be expected to develop 2 additional interactions. Similarly, eq. (6) shows that when *S* is large, we should expect a connectance which is a constant. Thus *p* has an interesting ecological interpretation: it represents the average connectance of networks large enough that the proportion (*S* − 1)/*S*^2^ is negligible.

### Other uses for our model

Our model is generative, and that is important and useful: we can use this model to correctly generate predictions that look like real data. This suggests that we can adapt the model, using eitherits parameters or predictions or both, to to get a new perspective on many questions in network ecology. Here we show four possible applications that we think are interesting, in that relying on our model eliminates the need to speculate on the structure of networks, or to introduce new hypotheses to account for it.

#### Probability distributions for *L_p_* and *Co*

In a beta-binomial distribution, it is assumed that the probability of success *p* varies among groups of trials according to a Beta(*μϕ*, (1 − *μ*)*ϕ*) distribution. Since *p* has a beta distribution, the linear transformations described by eq. (6) and eq. (7) also describe beta distributions which have been shifted and scaled according to the number of species *S* in a community. This shows that just as *L* must be within ecologically meaningful bounds, *Co* (eq. (6)) and *L_D_* (eq. (7)) must be as well. The connectance of a food web is bounded by (*S* − 1)/*S*^2^ and 1, while the linkage density is bounded by (*S* − 1)/*S* and *S*.

We can convert the beta distribution for *p* into one for *Co* by replacing *p* with the transformation of *Co* as described above (eq. (6)), and rescaling by the new range:

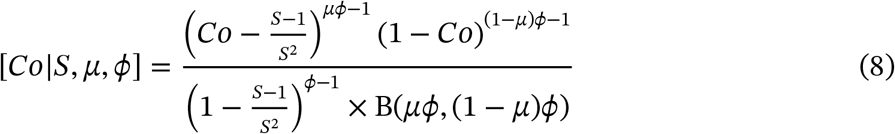

Similarly, we can convert the distribution for *p* into one for *L_D_* by replacing *p* with the transformation that gives *L_D_* (eq. (7))

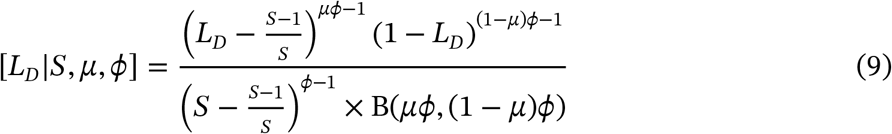

In fig. 3, we show that the connectance and linkage density obtained from the equations above fit the empirical data well.

**Figure 3:**
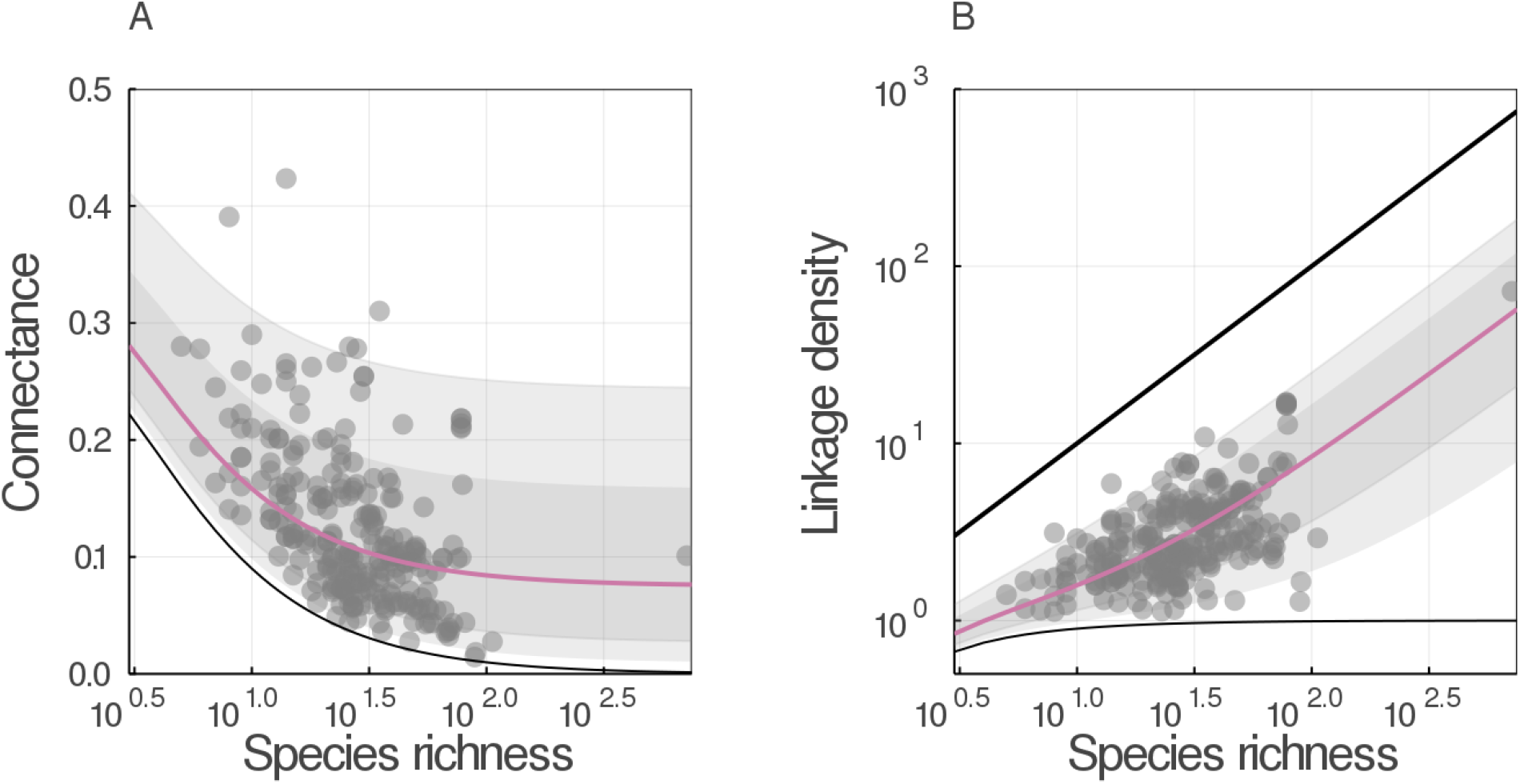
Connectance and linkage density can be derived from a model for links. A) Connectance and B) linkage density are plotted as a function of species richness, for the maximum *a posteriori* estimates of the flexible links model. In each panel, the colored line represent the median predicted quantity and the grey areas cover the 78% and 97% percentile intervals. Empirical data from the `mangal.io` database are plotted in each panel (grey dots). In A), the minimal (*S* − 1)/*S*^2^ connectance and in B) the minimal (*S* − 1)/*S* and maximum *S* linkage density are plotted (black lines).

#### An analytic alternative to null-model testing

Ecologists are often faced with the issue of comparing several networks. A common question is whether a given network has an “unusual” number of links relative to some expectation. Traditionally these comparisons have been done by simulating a “null” distribution of random matrices [21,22]. This is intended to allow ecologists to compare food webs to a sort of standard, hopefully devoid of whatever biological process could alter the number of links. Importantly, this approach assumes that (i) connectance is a fixed property of the network, ignoring any stochasticity, and (ii) the simulated network distribution is an accurate and unbiased description of the null distribution. Yet recent advances in the study of probabilistic ecological networks show that the existence of links, and connectance itself is best thought of as a probabilistic quantity [18]. Given that connectance drives most of the metrics of food web structure [17], it is critical to have a reliable means of measuring differences from the expectation. We provide a way to assess whether the number of links in a network (and therefore its connectance) is surprising. We do so using maths rather than simulations.

The shifted beta-binomial can be approximated by a normal distribution:

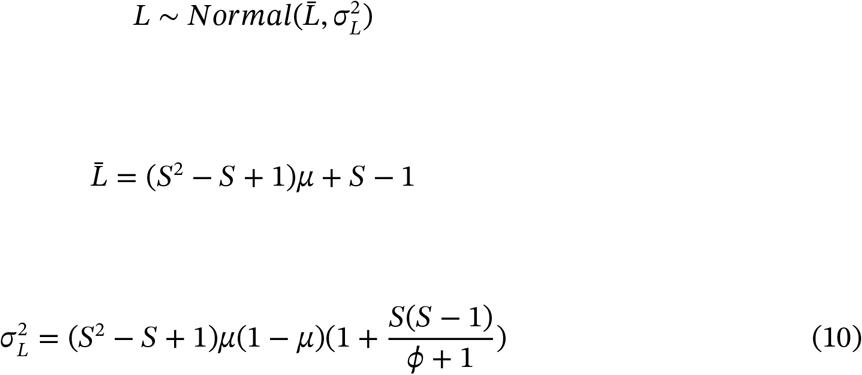

This means that given a network with observed species richness *S_obs_* and observed links *L_obs_*, we can calculate its *z*-score as

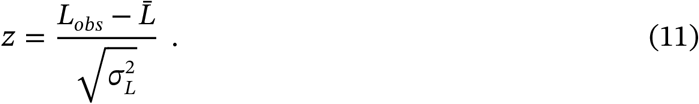

A network where 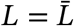 will have a *z*-score of 0, and any network with more (fewer) links will have a positive (negative) *z*-score. We suggest that the use of a *z*-score could help identify significantly under (over) sampled networks and estimate their number of missing (extra) links.

In fig. 4, we show that the predictions made by the normal approximation (panel B) are similar to those made by the beta distribution parameterized with the maximum *a posteriori* values of *μ* and *ϕ* (panel A), although the former can undershoot the constraint on the minimum number of links. This undershooting, however, will not influence any actual z-scores, since no food webs have fewer than *S* − 1 links and therefore no z-scores so low can ever be observed.

**Figure 4:**
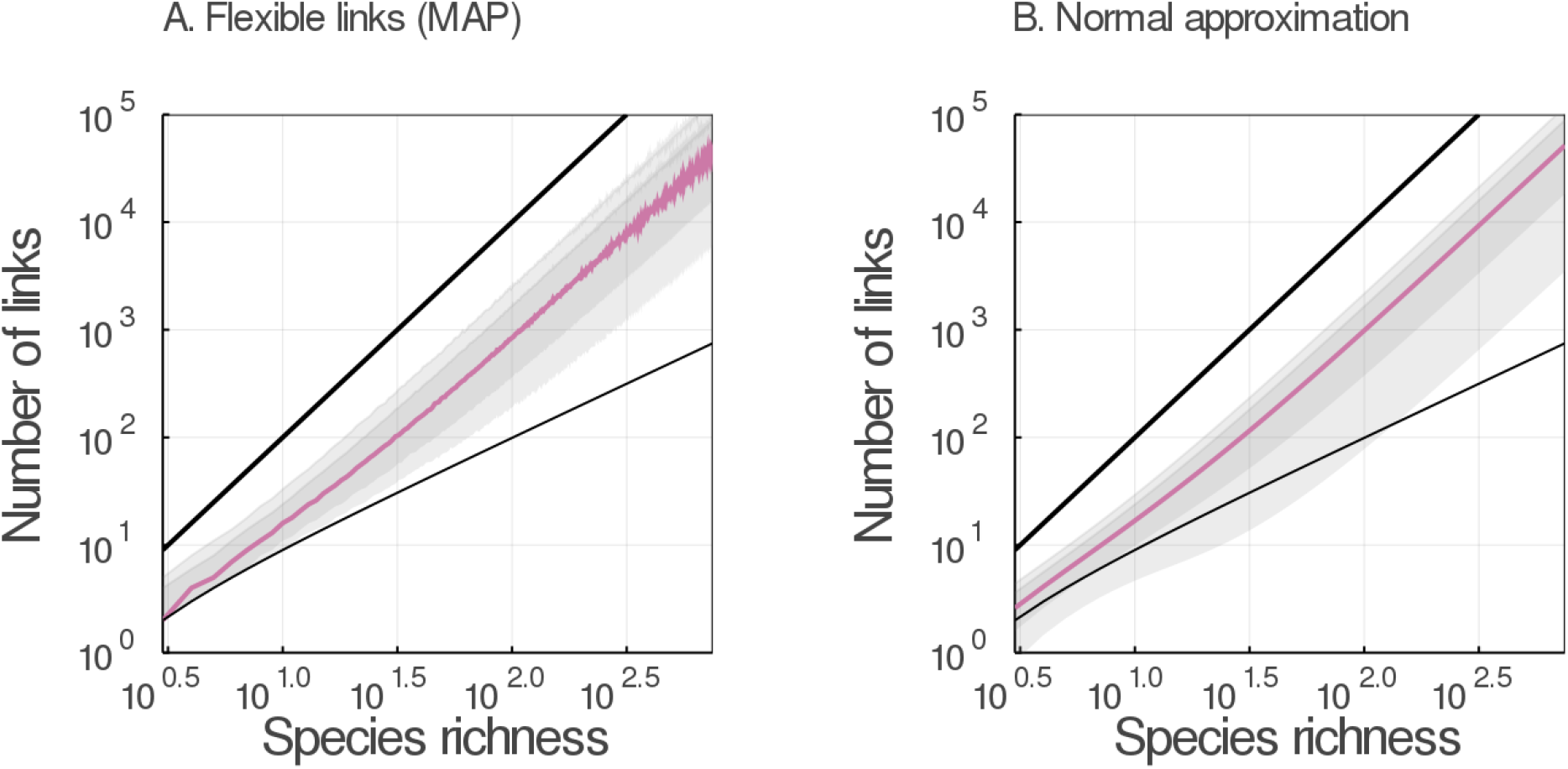
The shifted beta-binomial distribution can be approximated by a normal distribution. The number of links is plotted as a function of species richness obtained from A) the maximum *a posteriori* estimates of the flexible links model and B) its normal approximation. In each panel, the colored line represent the median predicted link number and the grey areas cover the 78% and 97% percentile intervals. The minimal *S* − 1 and maximal *S*^2^ numbers of links are plotted in each panel (thinner and bolder black lines, respectively)

#### We should see many different Network-Area Relationships

Our results bear important consequences for the nascent field of studying network-area relationships [23]. As it has long been observed that not all species in a food web diffuse equally through space [24], understanding how the shape of networks varies when the area increases is an important goal, and in fact underpins the development of a macroecological theory of food webs [25].

Using a power-law as the acceptable relationship between species and area [26,27], the core idea of studying NAR is to predict network structure as a consequence of the effect of spatial scale on species richness [23]. Drawing on these results, we provide in fig. 5 a simple illustration of the fact that, due to the dispersal of values of *L*, the relationship between *L/S* and area can have a really wide confidence interval. While our posterior predictions generally match the empirical results on this topic [28], they suggest that we will observe many relationships between network structure and space, and that picking out the signal of network-area relationships might be difficult.

**Figure 5:**
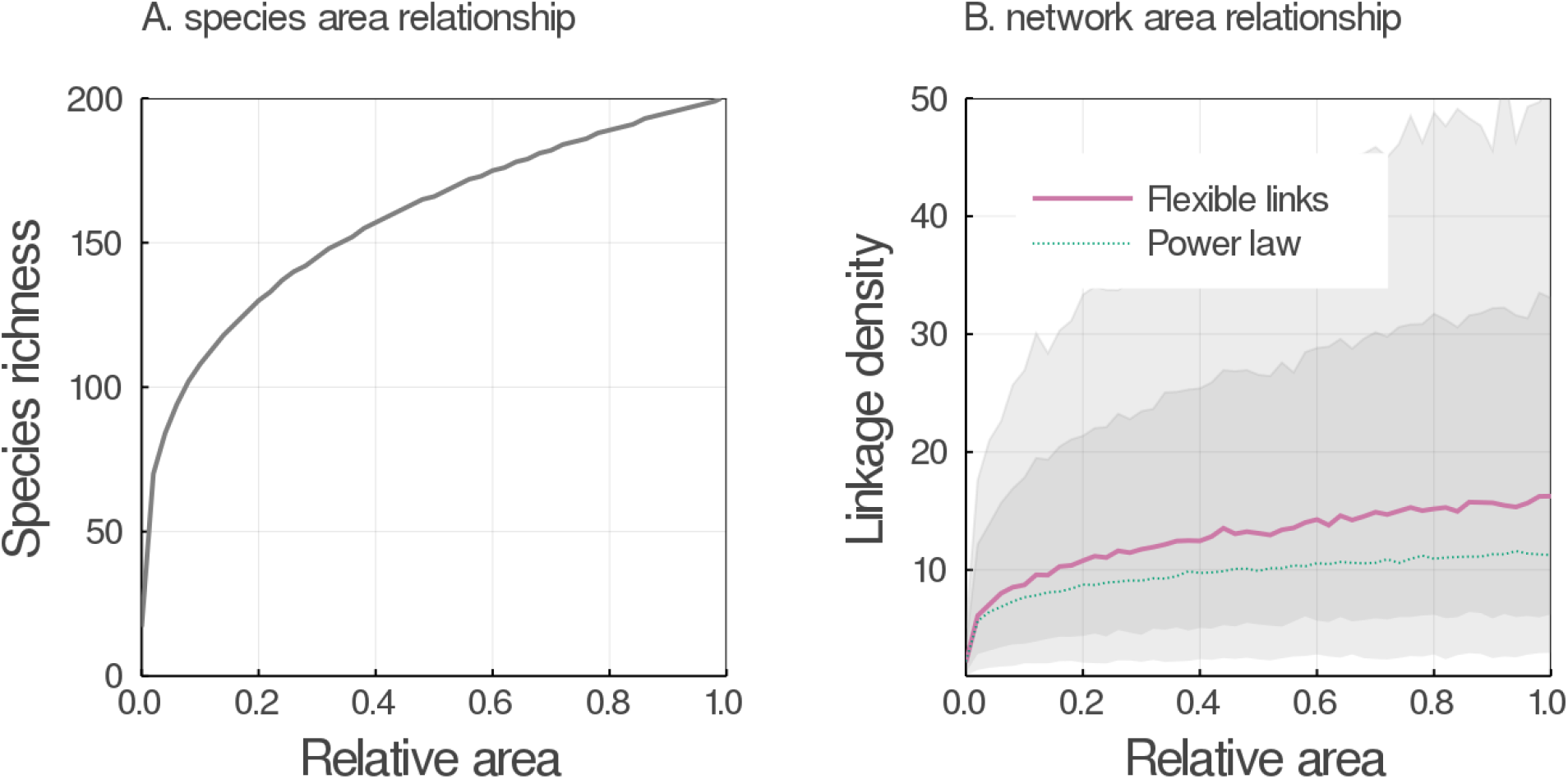
Many different Network-Area Relationships are supported by the data. Representing the species richness as *S* = *k* × *A^z^* (panel A), with *A* being the relative area size, *k* = 200 being the maximal species richness, and *z* = 0.27 a scaling exponent [23]. We then use the posterior distribution of *L* to predict how *L_D_* should scale with A. We compare the predictions of our model to that of the generally accepted powerlaw(eq. (3)). While our model predicts alargerlinkage density in larger areas (panel B), the confidence intervals around this prediction (grey areas covering the 78%and 97% percentile intervals) are extremely large. In particular, our model scales faster than the power law, but the confidence interval is high (due to the scaling of variance with *S*, eq. (10)). This suggests that we may observe either very weak, or very strong, effects of area on networks.

#### Stability imposes a limit on network size

Our model introduces a puzzling question: can organisms really interact with an infinite number of partners? According to eq. (7), at large values of *S*, the linkage density scales according to *p* × *S* (which is supported by empirical data), and so species are expected to have on average 2 × *p* × *S* interactions. A useful concept in evolutionary biology is the “Darwinian demon” [29], *i.e*. an organism that would have infinite fitness in infinite environments. Our model seems to predict the emergence of what we call Eltonian demons, which can have arbitrarily large number of interactions. Yet we know that constraints on handling time of prey, for example, imposes hard limits on diet breadth [30]. This result suggests that there are other limitations to the size of food webs; indeed, the fact that *L/S* increases to worryingly large values only matters if ecological processes allow *S* to be large enough. It is known that food webs can get as high as energy transfer allows [5], and as wide as competition allows [31]. In short, and as fig. 2 suggests, since food webs are likely to be constrained to remain within an acceptable richness, we have no reason to anticipate that *p* × *S* will keep growing infinitely.

Network structure may itself prevent *S* from becoming large. May [32] suggested that a network of richness *S* and connectance *Co* is stable as long as the criteria 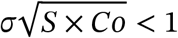 is satisfied, with *σ* being the standard deviation of the strengths of interactions. Although this criteria is not necessarily stringent enough for the stability of food webs [33,34], it still defines an approximate maximum value *σ*^⋆^ which is the value of above which the system is expected to be unstable. This threshold is 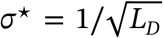, where *L_D_* is defined as in eq. (7). We illustrate this result in fig. 6, which reveals that *σ*^⋆^ falls towards 0 for larger species richness. The result in fig. 6 is in agreement with previous simulations, placing the threshold for stability at about 1200 species in food webs. These results show how ecological limitations, for example on connectance and the resulting stability of the system, can limit the size of food webs [33,35]. In the second panel, we show that networks of increasing richness (thicker lines, varying on a log-scale from 10^1^ to 10^3^) have a lower probability of being stable, based on the proportion of stable networks in our posterior samples.

**Figure 6:**
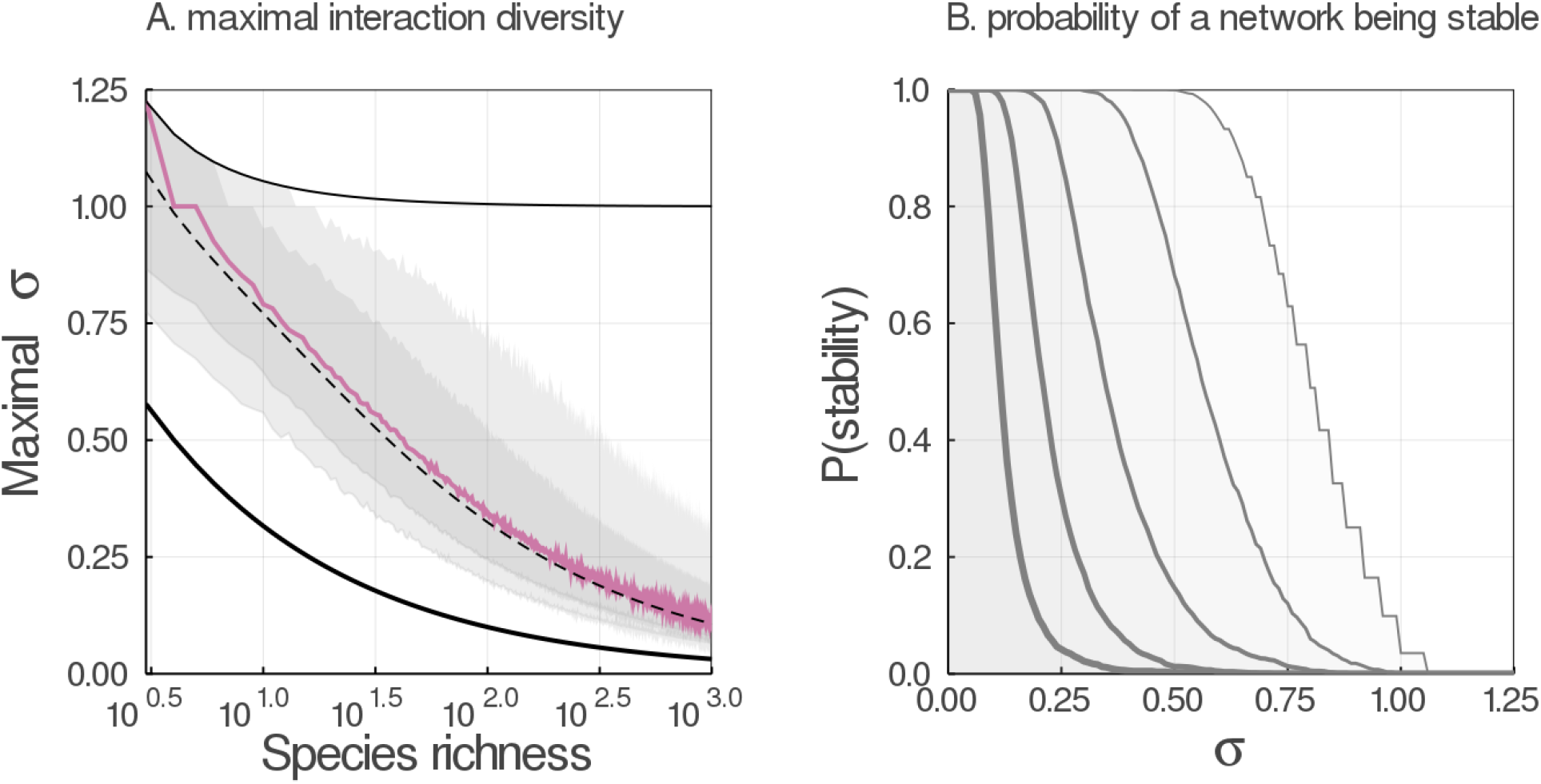
Stability imposes a limit on network size. Using eq. (7), we can calculate the maximum standard deviation in the strength of interactions which should ensure food web stability, 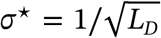 (panel A). The colored line represent the median value of maximum standard deviation, based on the posterior distribution of the flexible links model, and the grey areas cover the 78% and 97% percentile intervals. The fine and dark lines indicate the maximum and minimum value of maximum standard deviation, respectively. The dotted line shows the maximum for the average *L_D_*, as given by eq. (7). The maximum standard deviation falls sharply when the number of species increases, which will limit the stability of large food webs, and therefore explain why Eltonian demons should not emerge. In panel B, we show the probability of a network with *S* species being stable, based on draws from the posterior distribution, for 10 ≤ *S* ≤ 1000 - larger networks (thicker lines) are increasingly unlikely to be stable.

### Conclusions

Here we derived eq. (4), a model for the prediction of the number of links in ecological networks using a beta-binomial distribution for *L*, and show how it outperforms previous and more commonly used models describing this relationship. More importantly, we showed that our model has parameters with a clear ecological interpretation (specifically, the value of *p* in eq. (4) is the expected value of the connectance when *S* is large), and makes predictions which remain within biological boundaries.

This model also casts new light on previous results on the structure of food webs: small and large food webs behave differently [15]. Specifically, ecological networks most strongly deviate from scale free expectations when connectance is high [36]. In our model, this behaviour emerges naturally: connectance increases sharply as species richness decreases (fig. 3) - that is, where the additive term (*S* − 1)/*S*^2^ in eq. (6) becomes progressively larger. In a sense, small ecological networks are different only due to the low values of *S*. Small networks have only a very limited number of flexible links, and this drives connectance to be larger. Connectance in turn has implications for many ecological properties. Connectance is more than the proportion of realized interactions. It has been associated with some of the most commonly used network metrics [17], and contains meaningful information on the stability [36,37] and dynamics [38] of ecological communities. A probability distribution for connectance not only accounts for the variability between networks, but can be used to describe fundamental properties of food webs and to identify ecological and evolutionary mechanisms shaping communities. A recent research direction has been to reveal its impact on resistance to invasion: denser networks with a higher connectance are comparatively more difficult to invade [39]; different levels of connectance are also associated with different combinations of primary producers, consumers, and apex predators [40], which in turns determines which kind of species will have more success invading the network [41]. Because we can infer connectance from the richness of a community, our model also ties the invasion resistance of a network to its species richness.

The relationship between *L* and *S* has underpinned most of the literature on food web structure since the 1980s. Additional generations of data have allowed us to progress from the link-species scaling law, to constant connectance, to more general formulations based on a power law. Our model breaks with this tradition of iterating over the same family of relationships, and instead draws from our knowledge of ecological processes, and from novel tools in probabilistic programming. As a result, we provide predictions of the number of links which are closer to empirical data, stimulate new ecological insights, and can be safely assumed to always fall within realistic values. The results presented in fig. 5 (which reproduces results from [23]) and fig. 6 (which reproduces results from [33]) may seem largely confirmatory; in fact, the ability of our model to reach the conclusions of previous milestone studies in food web ecology is a strong confirmation of its validity. We would like to point out that these approaches would usually require ecologists to make inferences not only on the parameters of interests, but also on the properties of a network for a given species richness. In contrast, our model allows a real economy of parameters and offers ecologists the ability to get several key elements of network structure for free if only the species richness is known.

## Experimental Procedures

### Availability of code and data

All code and data to reproduce this article is available at the Open Science Framework (DOI: 10.17605/OSF.IO/YGPZ2).

### Bayesian model definitions

Generative models are flexible and powerful tools for understanding and predicting natural phenomena. These models aim to create simulated data with the same properties as observations. Creating such a model involves two key components: a mathematical expression which represents the ecological process being studied, and a distribution which represents our observations of this process. Both of these components can capture our ecological understanding of a system, including any constraints on the quantities studied.

Bayesian models are a common set of generative models, frequently used to study ecological systems. Here, we define Bayesian models for all 4 of the models described in eq. (1), eq. (2), eq. (3) and eq. (4). We use notation from [42], writing out both the likelihood and the prior as a product over all 255 food webs in the `mangal.io` database.

Link-species scaling (LSSL) model:

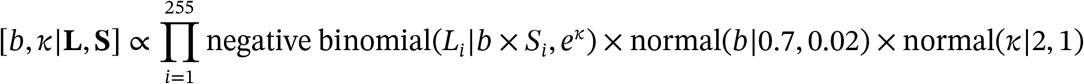

Constant connectance model:

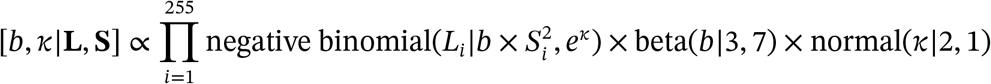

Power law model:

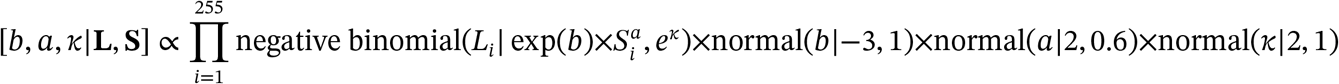

Flexible links model:

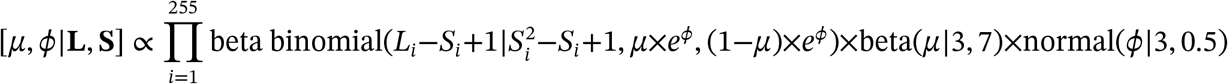

Note that while *e^ϕ^* is shown in these equations for clarity, in the text we use *ϕ* to refer to the parameter after exponentiation. In the above equations, bold type indicates a *vector* of values; we use capital letters for **L** and **S** for consistency with the main text.

Because we want to compare all our models using information criteria, we were required to use a discrete likelihood to fit all models. Our model uses a discrete likelihood by default, but the previous three models (LSSL, constant connectance and the power law) normally do not. Instead, these models have typically been fit with Gaussian likelihoods, sometimes after log-transforming *L* and *S*. For example, eq. (3) becomes a linear relationship between log(*L*) and log(*S*). This ensures that predictions of *L* are always positive, but allows otherwise unconstrained variation on both sides of the mean. To keep this same spirit, wechosethenegativebinomialdistributionfor observations. This distribution is limited to positive integers, and can vary on both sides of the mean relationship.

We selected priors for our bayesian models using a combination of literature and domain expertise. For example, we chose our prior distribution for *p* based on [12], who gave a value of constant connectance equal to 0.14. While the prior we use is “informative”, it is weakly so; as [12] did not provide an estimate of the variance for his value we chose a relatively large variation around that mean. However, no information is available in the literature to inform a choice of prior for concentration parameters *κ* and *ϕ*. For these values, we followed the advice of [43] and performed prior predictive checks. Specifically, we chose priors that generated a wide range of values for *L_i_*, but which did not frequently predict webs of either maximum or minimum connectance, neither of which are observed in nature.

### Explanation of shifted beta-binomial distribution

Equation eq. (4) implies that *L_FL_* has a binomial distribution, with *S*^2^ − *S* + 1 trials and a probability *p* of any flexible link being realized:

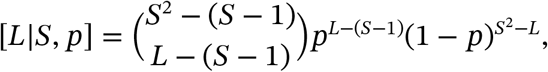

This is often termed a *shifted binomial distribution*.

We also note that ecological communities are different in many ways besides their number of species (*S*). Although we assume *p* to be fixed within one community, the precise value of *p* will change from one community to another. With this assumption, our likelihood becomes a shifted beta-binomial distribution:

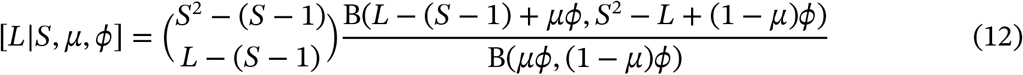

Where *B* is the beta function. Thus, the problem of fitting this model becomes one of estimating the parameters of this univariate probability distribution.

### Model fitting - data and software

We evaluated our model against 255 empirical foodwebs, available in the online database `mangal.io` [19]. We queried metadata (number of nodes and number of links) for all networks, and considered as food webs all networks having interactions of predation and herbivory. We use Stan [44] which implements Bayesian inference using Hamiltonian Monte Carlo. We ran all models using four chains and 2000 iterations per chain. Stan provides a number of diagnostics for samples from the posterior distribution, including 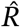, effective sample size, and measures of effective tree depth and divergent iterations. None of these indicated problems with the posterior sampling. All models converged with no warnings; this indicates that is it safe to make inferences about the parameter estimates and to compare the models. However, the calculation of PSIS-LOO for the LSSL model warned of problematic values of the Pareto-k diagnostic statistic. This indicates that the model is heavily influenced by large values. Additionally, we had to drop the largest observation (> 50 000 links) from all datasets in order to calculate PSIS-LOO for the LSSL model. Taken together, this suggests that the LSSL model is insufficiently flexible to accurately reproduce the data.

